# Impact of acute complex exercise on inhibitory control and brain activation: a functional near-infrared spectroscopy study

**DOI:** 10.1101/2022.10.02.510532

**Authors:** Shinji Takahashi, Philip M. Grove

## Abstract

A growing body of literature demonstrates that a single bout of exercise benefits executive function. While the acute effect of simple exercises like walking, running, and cycling has been well investigated, it is less clear how complex exercise, which requires open skills and various motions, impacts executive function and brain activation. Therefore, we compared the acute effects of a complex exercise on inhibitory control and brain activation with those of a simple exercise using functional near-infrared spectroscopy (fNIRS). Twenty-four young right-hand dominant adults (9 women) completed three interventions: badminton, running, and a seated rest control condition for 10 min each. During each intervention, oxygen uptake and heart rate were monitored. A Stroop task composed of neutral and incongruent conditions was administrated before and after each intervention. An fNIRS system recorded hemodynamics in the prefrontal cortex to evaluate brain activation during the Stroop task. The intensities of badminton and running were comparable. Performance on the Stroop task was significantly improved after badminton, specifically in the incongruent condition relative to in the neutral condition. On the other hand, neither running nor seated rest affected performance in the Stroop task. The fNIRS measures indicated that badminton and running had no significant influence on brain activation. These results show that a single bout of complex exercise enhances inhibitory control without increasing brain activation compared to simple exercise, suggesting that an acute complex exercise induces neural efficiency.

## Introduction

Regular exercise is known to play an essential role in both preventing or slowing cognitive decline in the elderly [1, 2] and enhancing the development of cognitive functions for children and adolescents[3–5] with several studies illuminating the association and mechanism between exercise and cognition. Investigations on the effect of a single bout of exercise on cognitive functions [6–9] have revealed effects on executive functions that include inhibitory control, working memory, and cognitive flexibility[10, 11]. So far, the accumulated evidence indicates that single bouts of aerobic exercise have a small but positive effect [5–8, 12], modulated by exercise intensity, duration, the timing of measures for executive functions[12], developmental stage[7], and baseline executive functions levels for individuals[6].

Although quantitative exercise features such as intensity and duration are well studied, qualitative features like exercise types have received less attention. In particular, the impact of acute complex exercises requiring open skills, adaptation to dynamic environments, and social interaction between teammates and opponents on executive functions is poorly understood. Regarding regular complex exercises, several systematic reviews indicate that regular complex exercises (e.g., basketball, football, and tennis) benefit executive functions more than simple exercises, such as closed skill sports (e.g., running and swimming)[3, 11, 13, 14]. Voss et al. [13] found that processing speed, a measure of cognitive functions, benefits from interceptive sports (e.g., tennis, squash) and strategic sports (e.g., volleyball, basketball, soccer). In contrast, static sports like long-distance running and swimming do not significantly affect processing speed. Contreras-Osorio et al.[3] demonstrated that sports interventions that involve complex exercises over several days have significant effects on the overall executive functions of children and teenagers. Despite contrary evidence, Diamond and Ling[11] argue that a simple exercise without a cognitive domain improves executive functions less than those with a cognitive domain, suggesting that an exercise with cognitive challenges and social components can enhance executive functions at every age.

Given the benefits of regular complex exercises on executive functions, it is expected that a complex exercise has different impacts on executive functions than a simple exercise for even single bouts of exercise. There have been several studies that investigated the various effects of acute exercise types on executive functions; however, the previous results are mixed[15–23].

Pesce et al.[17] reported that a team-based activity involving cognitive demands and social interaction enhanced free-recall word memory performance immediately after the activity. In contrast, a circuit activity with fewer cognitive demands and social interaction failed to show a similar effect. Cooper et al. [23] also demonstrated a significant impact of the team-based activity on inhibitory control and working memory. They found these processes were enhanced following basketball relative to seated rest. Similarly, Takahashi and Grove[18] found that singles-game badminton enhanced inhibitory control compared with seated rest. On the other hand, running on a motor-driven treadmill failed to enhance inhibitory control. These results support the idea that a single bout of complex exercise benefits executive functions relative to simple exercise, suggesting that open skills, various motions, and social interaction in complex exercise benefit executive functions and brain regions associated with them (e.g., prefrontal cortex, anterior cingulate cortex).

In contrast, other studies have reported that the effect of a single bout of complex exercise on executive functions is comparable to or less than that of simple exercises. O'Leary et al.[21] found that although walking on a motor-driven treadmill enhanced the inhibitory control and increased the amplitude of the P3 component, which reflects attentional resource allocation during cognitive tasks, exergaming composed of aerobic and cognitive domains did not positively affect the inhibitory control nor the amplitude of the P3 component. O’Leary et al. speculated that the aerobic nature of the exergaming enhances inhibitory control, however, the increased cognitive demands of exergaming offset the effect, resulting in no observed enhancement of the inhibitory control. Gallotta et al.[20] also failed to show a benefit of complex exercise on cognitive functions. Gallotta et al. compared basketball and running with seated rest, revealing that running and seated rest enhanced attention and concentration more than basketball. Like O’Leary, Gallotta et al. argued that the decision-making and the strategic manner required in basketball was an additional cognitive load that yielded mental fatigue, resulting in attenuating benefits of aerobic nature on attention and concentration. These results suggest that the cognitive demands of a single bout of complex exercise introduce additional burdens leading to mental fatigue canceling the beneficial effects of the aerobic nature on the executive functions. The above speculations are supported by previous studies[19, 24] showing that adding a cognitive task to a simple exercise increases cognitive burden compared with only a simple exercise and a cognitive task. The cognitive demand or mental fatigue could cancel out benefits from the aerobic nature to executive functions.

Although different impacts of complex exercises have been shown, studies reporting both positive and negative effects imply that a single bout of complex exercise would activate the brain more than simple exercises. Visser et al. [25] reported an interesting result regarding brain activation during a complex exercise. Visser et al. compared brain activation measured by electroencephalography during table tennis with activation during cycling, revealing increased brain activation in the frontal region during table tennis relative to cycling. Visser et al. demonstrated greater activation in the region of the brain underpinning the executive functions during a complex exercise relative to during a simple exercise. However, to the best of our knowledge, no study has compared the impact of simple and complex exercise on brain activation during an executive function task following the acute bout. Therefore, we measured brain activation during an inhibitory control task after a single session of simple exercise, complex exercise, and a rest control.

We compared behavioral performance (reaction time and accuracy) and hemodynamics in the prefrontal cortex after a bout of badminton, running, or seated rest in a cross-over experimental design, using a color-word Stroop task[26, 27] and functional near-infrared spectroscopy (fNIRS). Badminton was chosen as a complex exercise in this study because badminton requires various motions and cognitive demands. Running was the simple exercise. Given previous reports that a single aerobic exercise session improves behavioral measures of the Stroop task with increased hemodynamics in the prefrontal cortex [28–34], we hypothesized that a complex exercise like badminton would shorten reaction times in the Stroop task and increase hemodynamics in the prefrontal cortex compared to running.

## Materials and methods

### Participants

We conducted a power analysis for a one-way repeated analysis of variance (ANOVA) to calculate a sufficient sample size under the following conditions: the dependent variable was the change of the reaction time for the incongruent condition Stroop trial between pre-intervention and post-intervention; the independent variable was exercise mode three levels with badminton, running, and control intervention; partial eta squared was 0.05; power (1 - β) was 0.95; the intraclass correlation coefficient was 0.8[35] and alpha at 0.05. This analysis indicated that the sample size of 22 was adequate. Participants were recruited through sports and health sciences courses at Tohoku Gakuin University. The recruitment criteria were 1) right-hand dominant undergraduate students, 2) normal or corrected to normal vision, and 3) no history of brain, cognition, mental, or cardiovascular diseases. Twenty-four healthy right-hand dominant Japanese (9 women) voluntarily participated in this study. The participants were asked to refrain from alcohol use and strenuous physical activity for 24 h before each experiment and to refrain from smoking, food, or caffeine consumption for two hours preceding the experiments. Written informed consent was obtained from all participants before the first experiment. The Human Subjects Committee of Tohoku Gakuin University approved the study protocol (Approval number: 2019R003). Table 1 shows the characteristics of the participants.

**Table 1.** Characteristics of the participants (mean ± SE).

| Variable | Total ($N = 24$) | Men ($N = 15$) | Women ($N = 9$) |
| --- | --- | --- | --- |
| Age (years) | 20.4 ± 0.2 | 20.5 ± 0.1 | 20.2 ± 0.4 |
| Height (cm) | 168.2 ± 1.7 | 171.9 ± 1.8 | 161.9 ± 1.8 |
| Weight (kg) | 61.6 ± 1.8 | 66.1 ± 2.0 | 54.1 ± 1.9 |
| BMI ($\text{kg} \cdot \text{m}^{-2}$) | 21.7 ± 0.4 | 22.3 ± 0.5 | 20.6 ± 0.5 |
| IPAQ (MVPA-min · week$^{-1}$) | 2719.7 ± 652.2 | 3758.0 ± 937.1 | 989.3 ± 318.2 |
| VO<sub>2</sub>peak ($\text{mL} \cdot \text{kg}^{-1} \cdot \text{min}^{-1}$) | 46.9 ± 1.1 | 49.7 ± 1.1 | 42.1 ± 1.5 |
| HRpeak (bpm) | 192.9 ± 1.8 | 191.8 ± 2.5 | 194.8 ± 2.4 |

BMI: body mass index; IPAQ: International Physical Activity Questionnaire; MVPA: moderate-vigorous physical activity; VO<sub>2</sub>peak: the peak of oxygen uptake; HRpeak: the peak of heart rate.

### Procedures

Participants visited the sports physiology laboratory in the gymnasium on four different days (average interval, 6.1 ± 1.8 days).

#### Day 1

Participants received a brief introduction to the study and completed informed consent. Their height and weight were measured using a stadiometer and a digital scale. They then answered a questionnaire about medical history and the International Physical Activity Questionnaire (IPAQ)[36, 37], a measurement tool to assess the quantity of physical activity for the recent week. Next, to familiarize participants with the computer-based color-word Stroop task and an fNIRS device (OEG-16; Spectratech Inc., Yokohama, Japan), they completed the Stroop task while wearing the fNIRS device. A graded exercise test was subsequently conducted to determine the peak of oxygen uptake (VO_2_peak) and the peak of heart rate (HRpeak). After the graded exercise test, the participants completed the Stroop task with the fNIRS device again.

#### Day 2-4 (experimental sessions)

Laboratory visits 2, 3, and 4 were experimental sessions. To minimize any order or learning effects, the orders of the experimental sessions were randomized. After arrival at the laboratory, the participants rested on a comfortable chair for 10 min. In the experimental sessions, first, participants were asked about their mood by the two-dimensional mood scale (TDMS) [38] for 1 min to evaluate mental fatigue. They then completed the Stroop task with the sensor of fNIRS on their forehead before and after each intervention. Fitting the fNIRS sensor and calibrating this device took approximately 3 min, and the Stroop task took 4.2 min. After the pre-test of the Stroop task, participants removed the fNIRS sensor and were equipped with a portable indirect calorimetry system (MetaMax-3B; Cortex, Leipzig, Germany) in 1 min. Participants then rested on a chair for 3 min. For the badminton intervention, the participants moved from the laboratory to a badminton court, which took 2 min. For both the running and the control interventions, the participants walked on a treadmill (O2road, Takei Sci. Instruments Co., Niigata, Japan) at 4.2 km·h^−1^ for 2 min, which served as a counterpart to the move from the laboratory to the badminton court. Subsequently, the participants performed each intervention. Based on the protocol of the previous studies[16, 18, 35], the duration of the intervention was set to 10 min. After each intervention, participants returned to the laboratory or walked on the treadmill for 2 min and then they removed the indirect calorimetry system. Subsequently, they rested for 15 min on the chair. At the end of the resting period, the participants’ moods were probed by the TDMS again. Finally, they completed the post-test of the Stroop task and the fNIRS measure again.

In the badminton intervention, the participants played a singles game against an investigator who had experience teaching badminton in physical education courses. The investigator played at a level of proficiency that matched the participant’s level and had social interaction with participants by providing tips. During the game, the scores were not recorded, and “victory or defeat” was not determined. In the running intervention, the participants ran on the treadmill. Running speed was set according to each participant’s 75%VO_2_peak, which has been previously shown to be the intensity equal to that of the badminton intervention [18, 35]. In the control intervention, the participants were seated on a comfortable chair with their smartphones and were instructed to spend time operating their smartphones as normal. Oxygen uptake (VO_2_), carbon dioxide output (VCO_2_), and HR were monitored throughout each experimental session. Physiological measures for the last 7 min were averaged, and the rating of perceived exertion (RPE) was evaluated at the end of each intervention.

### Aerobic fitness assessment

Participants performed the graded exercise test on the motor-driven treadmill to volitional exhaustion. The initial speed was set at 7.2 km·h^−1^. The slope of the treadmill was constant at 1.0%. Each speed lasted 2 min and the speed was increased by 1.2 km·h^−1^ until volitional exhaustion. The MetaMax-3B measured VO_2_, VCO_2_, and HR, and the average of the final 30 s was defined as VO_2_peak and HRpeak. RPE was measured at the end of each stage. Volitional exhaustion was reached based on the following criterion: 1) RPE ≥ 17, 2) HR ≥ 95% of age-predicted HRmax (220 minus age), and 3) a respiratory exchange ratio (RER, VCO_2_·VO_2_^−1^) ≥ 1.10.

### Color-word Stroop task

The computer-based color-word Stroop task composed of a neutral condition block and an incongruent condition block assessed inhibitory control for each participant (see Fig 1). The beginning of the Stroop task was the baseline period of 30 s, in which a picture of nature (river scenery) was presented on a 15-inch screen of a laptop computer. After the baseline period, either the neutral condition block or the incongruent condition block was started. The order of the neutral and incongruent condition blocks was randomized. Each condition block had 24 trials. For the trials in the neutral condition, the laptop screen presented two rows on a gray background, the upper row was ‘XXXX’ printed in one of the red, blue, yellow, and green, and the lower row was one of the words ‘Red,’ ‘Blue,’ ‘Yellow,’ and ‘Green’ printed in white. For the trials in the incongruent condition, the upper row presented either word ‘Red,’ ‘Blue,’ ‘Yellow,’ and ‘Green’ printed in incongruent colors (e.g., ‘Red’ was printed in blue), and the lower row was one of the words ‘Red,’ ‘Blue,’ ‘Yellow,’ and ‘Green’ printed in white. The trials were shown on the screen for 3 s. The upper row was presented 500 ms before the lower row. Each trial for the neutral and incongruent conditions was separated by intervals that showed a white cross for 1 s.

**Fig 1.**
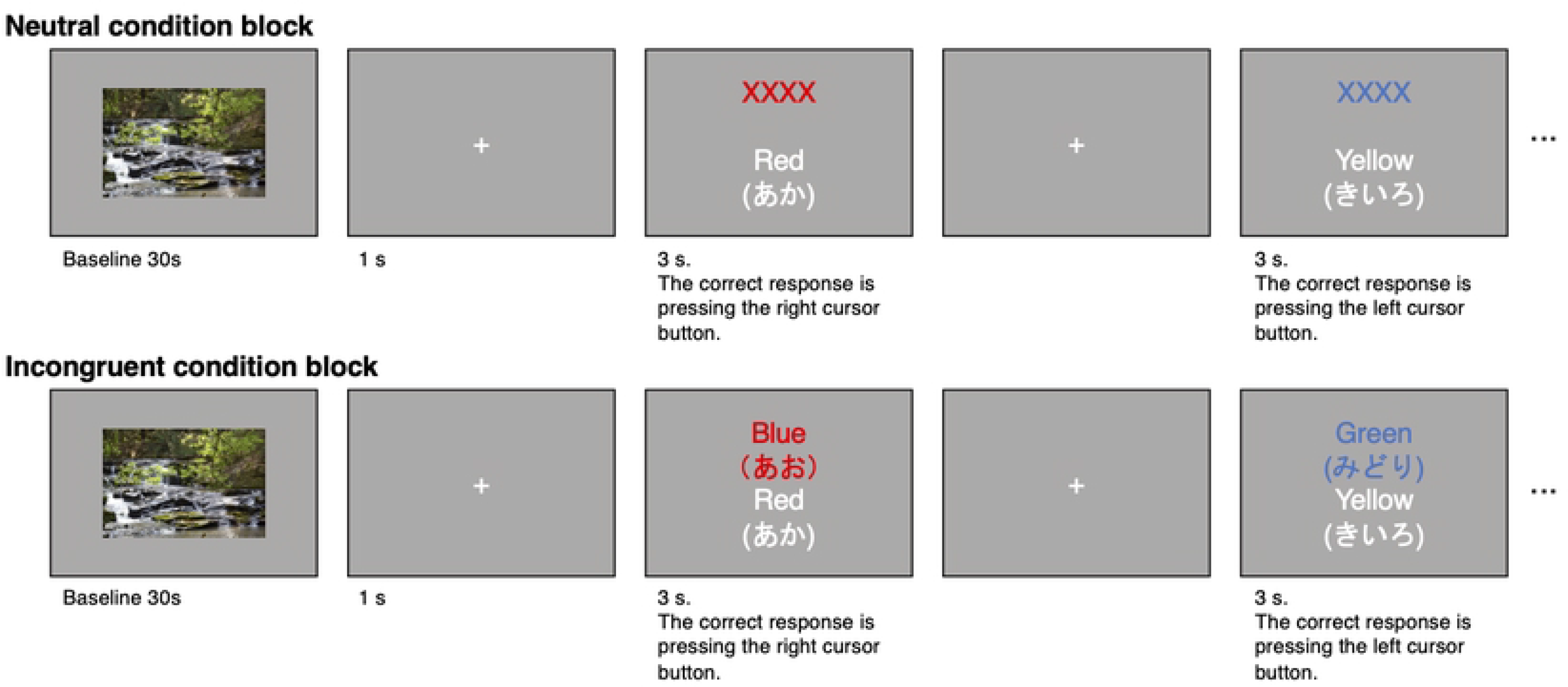
Scheme of the color-word Stroop task. The scheme consists of two blocks the neutral condition block and the incongruent condition block. The order of the two blocks was randomized.

The participants were instructed to press the right cursor button with the right hand when the color of the upper row matched the color name in the lower row and to press the left cursor button with the right hand when the color of the upper row was mismatched with the color name in the lower row. The ratio of the correct responses of the right (matched) and left (mismatched) buttons were set to 50%, and the order of the two answers was randomly assigned. The participants were asked to respond as quickly and accurately as possible. All words in the task were written in Japanese. Reaction time and accuracy response rate (%) were recorded.

### Functional near-infrared spectroscopy

The multi-channel fNIRS device (OEG-16) measured hemodynamics in the prefrontal cortex during the Stroop task. This device has a headband module with six emitter probes and six detector probes arranged on a two × six matrix to detect signals from 16 channels. The six emitter probes emit two-wavelength near-infrared light (approximately 770 and 840 nm). The six detector probes placed at 30 mm from the emitter probes detect the re-emitted light. The center of the probe matrix is placed on Fpz (International 10-20 system), and the matrix covers from F7 to F8 (see Fig 2). This device records changes in oxy-hemoglobin (oxy-Hb) and deoxy-hemoglobin (deoxy-Hb) at approximately 30 mm below the scalp. The sampling interval is 0.65 s. We cleaned the emitter, detector probes, and the forehead area for participants with alcohol cotton before each Stroop task.

**Fig 2.**
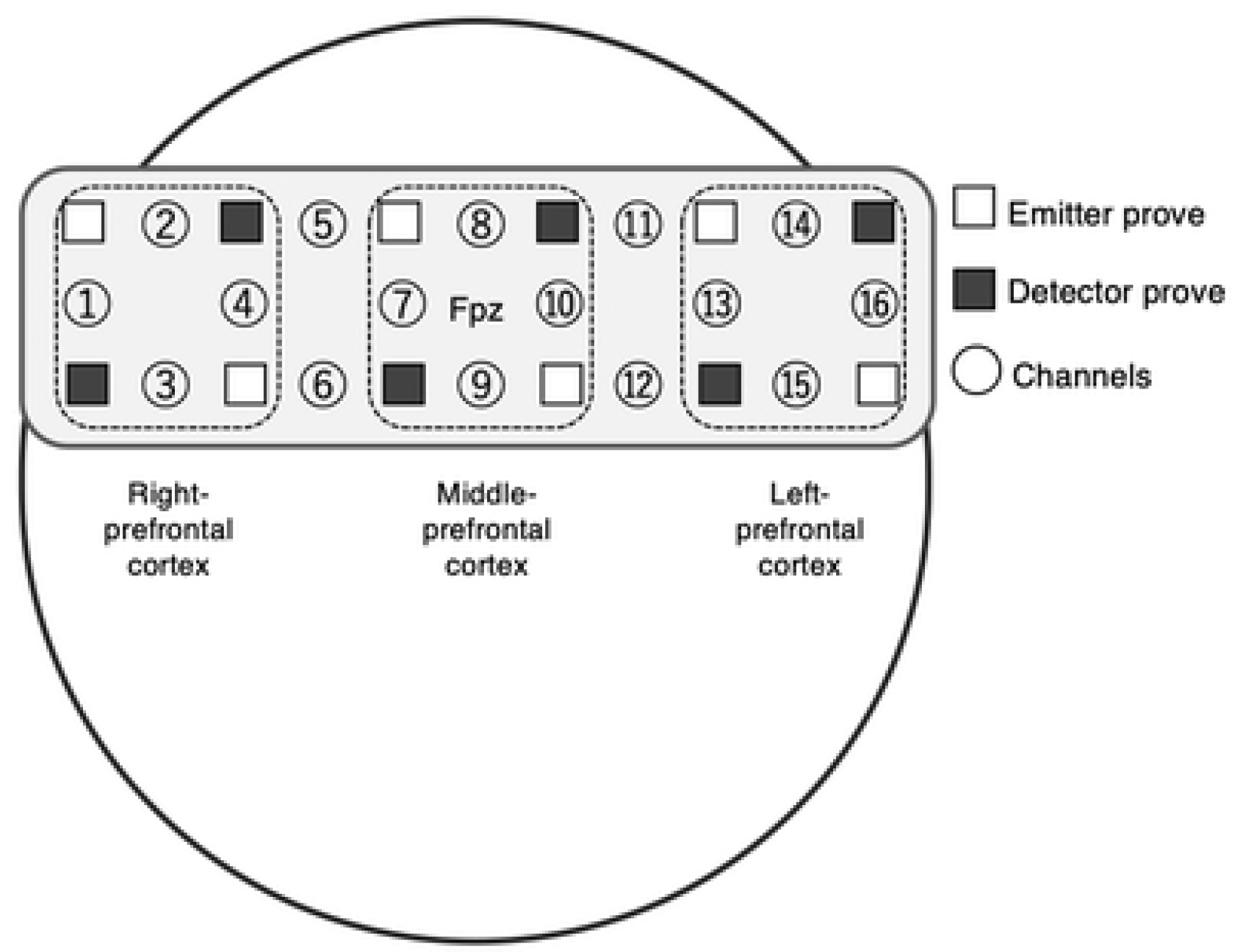
The layout for emitter and detector probes in the headband module of the NIRS device. The center of the probe matrix was placed on the Fpz (International 10– 20 System).

### Processing the fNIRS data

Raw records of oxy-Hb and deoxy-Hb were band-pass filtered on the specific software for this device (OEG-16 control program). To remove artifacts from drift, respiration, and heartbeat, the low-pass and high-pass filters were set at 0.005 Hz and 0.05 Hz, respectively. Moreover, artifacts from changes in skin blood flow in the forehead were removed using the hemodynamic modality separation method[39]. After this data processing, the dataset of raw data was exported to the CSV format file. Then, in order to increase the ratio of signal, oxy-HB and deoxy-Hb for each channel were converted to *z*-scores using the mean and standard deviation for the first 13 s (20 samples) of the first baseline in the Stroop task. We used data for the first 13 s of the first baseline for the calculation of *z*-scores because the participants appeared to mentally prepare for the Stroop task so that there was an increase and decrease trend in oxy-Hb and deoxy-Hb during the latter part of the baseline. Since the *z*-scores for oxy-Hb and deoxy-Hb modified by the hemodynamic modality separation method[39] resulted in an identical value, although opposite in sign, we used the oxy-Hb *z*-scores for subsequent analyses.

We averaged the oxy-Hb in every channel for the neutral and incongruent condition blocks. Then, the oxy-Hb *z*-scores for the neutral and incongruent condition blocks were averaged for 1-4 channels (Right-prefrontal cortex), 7-10 channels (Middle-prefrontal cortex), and 13-16 channels (Left-prefrontal cortex), respectively.

### Psychological mood

To assess the subjective mental fatigue of participants that potentially affects behavioral performance for the Stroop task, the TDMS[38] measured psychological pleasure and arousal states before the Stroop tasks. The TDMS was composed of eight items (6 Likert scales) that probed the psychological conditions of energetic, lively, lethargic, listless, relaxed, calm, irritated, and nervous. The range of pleasure and arousal states scored by the TDMS was −20 to 20.

### Statistical analysis

All measurements were described as mean ± standard error of the mean. Statistical analyses were conducted using IBM SPSS 27 (SPSS Inc., Chicago, IL, USA). To examine the exercise intensity of each intervention, %HRpeak, %VO_2_peak, RER, and RPE were compared using the mixed model with Mode (running, badminton, and control) and Order of interventions (moderator) as a fixed effect and participant as a random effect. A significant main effect of Mode was followed up with the Bonferroni corrected *t*-tests.

Psychological moods scored by the TDMS were analyzed using the mixed model with Time (pre-and post-test) and Mode (running, badminton, and control) and the interaction (Time × Mode) and Order of interventions (moderator) as fixed effects and participant as a random effect for pleasure level and arousal level, respectively. If there was a significant interaction, psychological moods were compared using the mixed model with Mode and Order of interventions (moderator) as a fixed effect and participant as a random effect and the Bonferroni corrected *t*-tests for pre-test and post-test, respectively.

The reaction time and accuracy response rate for the Stroop tasks were compared using the mixed model with Condition (neutral and incongruent), Time (pre- and post-test), and Mode (running, badminton, and control), interactions (Condition × Time, Condition × Mode, Time × Mode, and Condition × Time × Mode), Order of interventions (moderator), and Order of condition blocks for the Stroop task (moderator) as fixed effects and participant as a random effect. When any significant interactions were found, the mixed models with two fixed effects composing the significant interactions and the participant as a random effect were examined as a post hoc analysis.

Similarly, the oxy-Hb for fNIRS was analyzed using the mixed model with Condition, Time, Mode, interactions, Order of interventions (moderator), and Order of condition blocks for the Stroop task (moderator) as fixed effects and participant as a random effect, for the Right-, Middle-, and Left-prefrontal cortex, respectively.

We set the covariance structure as the unstructured for all the mixed models. Significance levels for all analyses were set at *P* = .05. Cohen’s *d* was calculated to assess the effect sizes for differences between the two means.

## Results

### Intensity of interventions

Table 2 presents the intensities for each intervention. The mixed models for %VO_2_peak, %HRpeak, RER, and RPE revealed the significant main effects (*F*s (2, 20.9-22.9) ≥ 15.4, *P*s < .001). The % VO_2_peak, %HRpeak, RER, and RPE during both the badminton and running interventions were significantly higher than during the control intervention (*P*s < .001, Cohen’s *d*s ≥ 1.14). Differences in all intensity measures between the badminton and running interventions were not significant (*P*s ≥ .084, Cohen’s *d*s ≤ |0.50|).

**Table 2.** Intensities of each intervention (mean ± SE).

| Variable | Intervention | Mean± SE (N = 24) |
| --- | --- | --- |
| %VO<sub>2</sub>peak (%) | Badminton | 77.9 ± 1.9 * |
| | Running | 76.9 ± 0.9 * |
| | Control | 9.2 ± 0.3 |
| %HRpeak (%) | Badminton | 83.3 ± 1.5 * |
| | Running | 87.3 ± 0.9 * |
| | Control | 38.0 ± 1.1 |
| RER (VCO<sub>2</sub>·VO<sub>2</sub><sup>-1</sup>) | Badminton | 0.96 ± 0.02 * |
| | Running | 0.98 ± 0.01 * |
| | Control | 0.89 ± 0.01 |
| RPE | Badminton | 12.6 ± 0.4 * |
| | Running | 13.6 ± 0.3 * |
| | Control | 6.2 ± 0.3 |

\* Significantly different from control intervention; $p < .05$ at Bonferroni multiple comparison tests.

### Psychological mood

Table 3 shows the changes in the pleasure and arousal state scored by the TDMS for each intervention. The mixed model for the pleasure state did not reveal significant main effects of Mode (*F* (2, 22.4) = 2.8, *P* = .083), Time (*F* (2, 23.0) = 0.5, *P* = .475) or the interaction of Mode × Time (*F* (2, 23.0) = 2.4, *P* = .112). The mixed model for the arousal state revealed a significant interaction of Mode × Time (*F* (2, 23.0) = 14.4, *P* < .001) and the main effects of Mode (*F* (2, 19.1) = 33.7, *P* < .001) and Time *F* (2, 23.0) = 34.3, *P* < .001). To decompose the significant interaction of Mode × Time for the arousal state, we compared the arousal state of each intervention for pre-test and post-test, respectively. While there was no significant main effect of Mode (*F* (2, 20.0) = 1.7, *P* = .206) for pre-test, the main effect of Mode was significant (*F* (2, 19.8) = 34.6, *P* < .001) for post-test. The arousal states after the badminton and running interventions were significantly higher than after the control intervention (*P*s < 0.001, Cohen’s *d*s ≥ 1.01). The difference between the badminton and the running interventions at post-test was not significant (*P* = .980, Cohen’s *d* = 0.23).

**Table 3.** Psychological states in each intervention (mean ± SE).

| Variable | Intervention | Pre-test | Post-test |
| --- | --- | --- | --- |
| Pleasure state<br>(arbitrary<br>unit) | Badminton | 8.7 ± 0.9 | 10.0 ± 0.8 |
| | Running | 8.3 ± 1.0 | 8.3 ± 0.8 |
| | Control | 8.2 ± 0.8 | 8.3 ± 1.0 |
| Arousal state<br>(arbitrary<br>unit) | Badminton | -3.5 ± 0.7 | 2.7 ± 0.7* |
| | Running | -3.7 ± 0.8 | 1.4 ± 1.0* |
| | Control | -4.6 ± 0.6 | -4.9 ± 0.9 |

\* Significantly different from control intervention for post-test; $p < .05$ at Bonferroni multiple comparison tests.

### Color-word Stroop task

The mixed model for the accuracy response rate of the color-word Stroop task revealed the significant main effect of Condition (*F* (1, 22.7) = 202.1, *P* < 0.001) but none of the other interactions or main effects were significant (*F*s (1-2, 21.5-23.0) ≤ 3.3, *P*s ≥ .057). The accuracy response rate for the neutral condition was significantly higher than for the incongruent condition (99.0 ± 0.3% vs. 94.8 ± 0.3%, *P* < .001, Cohen’s *d* = 2.98).

Fig 3 presents the changes in the reaction time for each Mode. The mixed model for the reaction time revealed a significant interaction of Mode × Time (*F* (2, 23.0) = 5.6, *P* = .011), a main effect of Time (*F* (1, 23.0) = 14.6, *P* < .001), and a main effect of Condition (*F* (1, 23.1) = 29.7, *P* < .001). The main effect of Mode (*F* (2, 21.1) = 0.3, *P* = .765) and the other interactions (*F*s (1-2, 23.0-23.2) ≤ 1.5, *P*s ≥ .250) were not significant. To decompose the significant interaction of Mode × Time, we analyzed reaction time using the mixed model with Time, Condition, Time × Condition, Order of interventions, and Order of conditions as the fixed effects and the random effect of the participants for each intervention.

**Fig 3.**
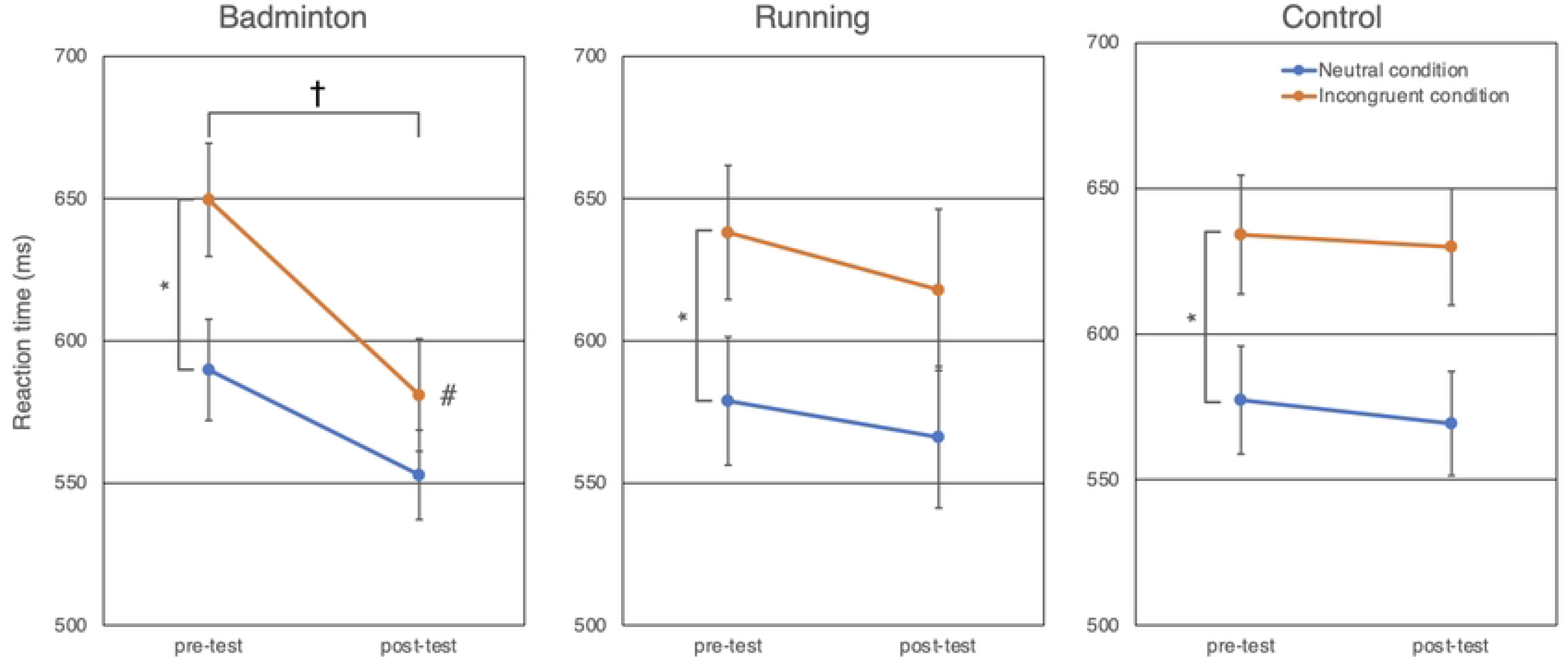
Comparisons of the reaction time for each intervention. Blue lines represent changes in the reaction time for the neutral condition. Red lines represent changes in the reaction time for the incongruent condition. Error bars show the standard errors of the mean. For the badminton intervention, the reaction time for both neutral and incongruent conditions was shortened, and the decrease of the reaction time for the incongruent condition was more prominent than for the neutral condition. Changes between pre-test and post-test were not significant for the running and control conditions. * means the significant difference between the neutral condition and the incongruent condition (*p* < .05), † means the significant difference between pre-test and post-test (*p* < .05), and # means that the reaction time for the incongruent condition was significantly shortened than that for the neutral condition (*p* < .05).

For the badminton intervention, the interaction of Time × Condition (*F* (1, 23.2) = 6.4, *P* = .018), the main effects of Time (*F* (1, 23.0) = 26.7, *P* < .001), and Condition (*F* (1, 23.0) = 14.5, *P* < .001) were all significant. The significant interaction showed that reaction times were shorter for badminton for the incongruent condition (Cohen’s *d* = −0.71) relative to that for the neutral condition (Cohen’s *d* = −0.48). For the running and control interventions, although the main effects of Condition were significant (*F*s (1, 22.3-22.9) ≥ 16.3, *Ps* < .001), the interactions of Condition × Time (*F*s (1, 22.3-23.7) ≤ 0.1, *Ps* ≥ .769) and the main effects of Time (*F*s (1, 23.0) ≤ 1.3, *Ps* ≥ .263) were not significant.

### Hemodynamics in the prefrontal cortex

Fig 4 presents the changes in oxy-Hb (*z*-score) during the Stroop task from the pre-test to post-test for the right-, middle-, and left-prefrontal cortex, respectively. Mixed model analysis did not reveal any significant main effects or interactions for the right-prefrontal cortex region (*F*s (1-2, 18.8-23.0) ≤ 2.7, *P*s ≥ .089). There were no significant main effects or interactions for the middle-prefrontal cortex region (*F*s (1-2, 20.7-23.0) ≤ 0.7, *P*s ≥ .444). A significant main effect of Condition (*F* (1, 23.4) = 7.0, *P* = .014) for the left-prefrontal cortex was found but no other main effects or interactions were significant (*F*s (1-2, 21.3-23.0) ≤ 1.2, *P*s ≥ .317). These results indicated that oxy-Hb levels in the left-prefrontal cortex were significantly increased in the incongruent condition relative to the neutral condition (the incongruent condition 5.4 ± 0.6 vs. the neutral condition 4.6 ± 0.4, Cohen’s *d* = 0.55), regardless of the timing relative to exercises or rest.

**Fig 4.**
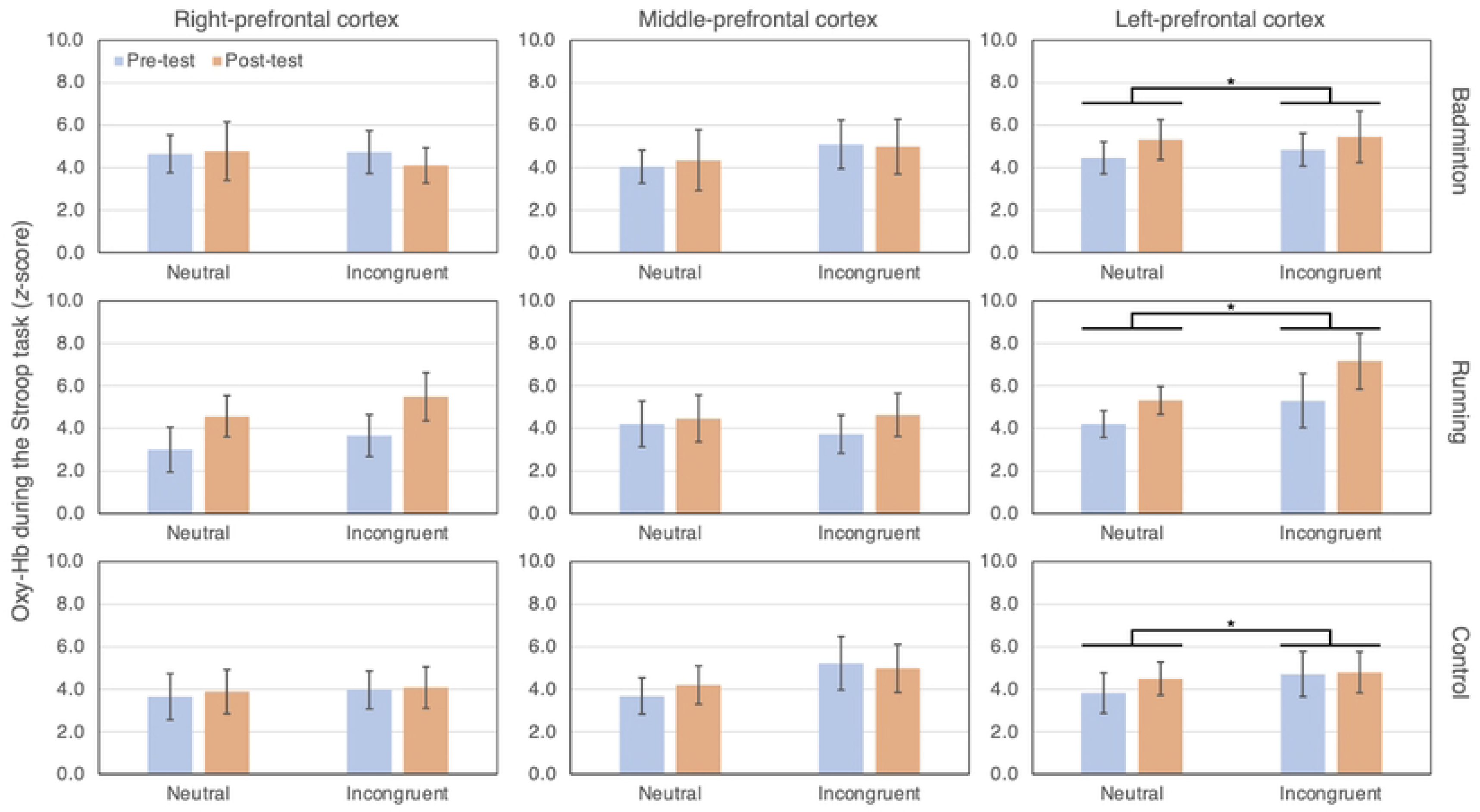
Comparisons of the oxygenated hemoglobin (z-score) for each intervention and region of the prefrontal cortex. Blue bars represent oxygenated hemoglobin at pre-test. Red bars represent oxygenated hemoglobin at post-test. Error bars show standard errors of the mean. * means the significant difference between the neutral and incongruent conditions (*p* < .05) for the left-prefrontal cortex throughout all interventions.

## Discussion

This study investigated the impact of a single bout of complex exercise on inhibitory control and brain activation compared to simple exercise. To the best of our knowledge, this study is the first to measure brain activation after a single bout of complex exercise. The main findings of this study were that a single bout of badminton yielded shorter reaction times for the neutral and incongruent conditions compared to simple exercise and a rest control while maintaining the same accuracy response rate. Furthermore, the acute effect of badminton on performance in the incongruent condition, requiring a greater amount of inhibitory control [40], was more prominent than in the neutral condition. On the other hand, no differences between badminton and running and between pre-test and post-test were observed in the oxy-Hb for each prefrontal cortex region. Taken together, our results indicate that a single bout of complex exercise enhances inhibitory control compared to simple exercise without increased hemodynamics in the prefrontal cortex underpinning the inhibitory control. This suggests that an acute complex exercise leads to more efficient functioning of the prefrontal cortex, enhancing executive functions relative to acute simplex exercise.

Both the badminton and running interventions were vigorous-intensity [41] exercises for a short duration of 10 min. The badminton intervention enhanced inhibitory control but this was not found after the running intervention. Our failure to observe enhanced inhibitory control following the running intervention with vigorous-intensity and duration of 10 min is consistent with previous studies. A previous meta-analysis [12] reported that a single session of aerobic exercises with vigorous-intensity has significantly smaller effects on cognitive functions relative to very-light, light, and moderate-intensity exercises. Further, Chang et al. [42] investigated the dose-response relationship between the duration of acute exercise and its effect on inhibitory control, revealing that the exercise time of 10 m was too short to increase performance of inhibitory control. It seems reasonable that a simple aerobic exercise with vigorous-intensity and short duration has no effect on inhibitory control.

Although the physiological parameters and exercise-induced changes in psychological moods were comparable during the badminton and running interventions, shorter reaction times for the neutral and incongruent conditions with maintaining the accuracy response rate were only observed in the badminton intervention. Additionally, the effect of badminton on the incongruent condition was more prominent than on the neutral condition. These results suggest that the features of badminton, such as open skills, various motions, and social interaction, are causes of more extent effects on inhibitory control, supporting our hypothesis. Unlike the Stroop task performance results, the fNIRS results were inconsistent with our hypothesis. Previous fNIRS studies [28–32] have indicated that the increased hemodynamics in the prefrontal cortex after a simple exercise can improve Stroop task performances. Given the previous results that the enhanced inhibitory control would be brought from the exercise-induced activated prefrontal cortex, we hypothesized that an acute complex exercise increases oxy-Hb during the Stroop task in the prefrontal cortex relative to a simple exercise, the increasing oxy-Hb improves inhibitory control. However, the shorter reaction times observed in the incongruent condition after badminton did not correlate with increasing oxy-Hb in all prefrontal cortex regions. Prefrontal cortex activation was comparable for both exercise interventions, but improvement in the Stroop task performance was only observed in the badminton intervention. These results suggest that a single bout of complex exercises could induce suitable conditions in the prefrontal cortex to achieve goals of inhibitory control tasks, resulting in a decreased Oxy-Hb level required for higher behavioral performance. That is, an acute complex exercise could induce neural efficiency [43–45] for executive functions.

Regarding neural efficiency following an acute vigorous-intensity and short-duration exercise, noteworthy findings were reported by Kao et al.[46]. Kao et al. compared 9 min of high-intensity interval training (HIIT) with 20 min of continuous aerobic exercise and 20 min of seated rest, finding that HIIT improved behavioral performances in a flanker task, although HIIT decreased the amplitude of P3, indicating lower attentional resource allocation relative to seated rest and continuous aerobic exercise. Further, the latency of P3 reflecting cognitive processing speed was significantly shortened after HIIT compared with seated rest. Still, no difference in the latency of P3 was observed between continuous aerobic exercise and seated rest. Kao et al.’s results indicate that there are different mechanisms in the acute effect of exercise on executive functions, and types of exercise modulate it.

Kao et al. speculated that exercise-induced increasing serum lactate could be a potential reason for neural efficiency after exercise. A vigorous-intensity exercise accumulates serum lactate, allowing the brain to uptake lactate as an energy source, in addition to glucose. Therefore, they proposed that such a shift in energy sources could induce neural efficiency. However, our results do not seem to support the lactate hypothesis. In this study, because RER (VCO_2_·VO_2_ ^−1^) correlated with lactate [47, 48] was not different for badminton and running as well as other exercise-intensity indices, serum lactate concentration during badminton was comparable with during running. Assuming potentially comparable lactate for badminton and running, neural efficiency after badminton might be induced from the features of complex exercises.

We speculate that an activated brain during an acute complex exercise could be a potential reason for neural efficiency after acute complex exercise. Visser et al. [25] compared the brain activation measured by electroencephalography during table tennis with during cycling, finding that the frontal and central regions of the brain were more activated during table tennis than during cycling even though the intensity during cycling (HR was 103.2±15.2 bpm) was higher than table tennis (HR was 91.3±14.0 bpm). Considering the previous results in Visser et al., the brain during badminton could also be more activated than during running. Increased brain activation during complex exercise could put the brain into a suitable condition to exert inhibitory control following cessation of exercise, requiring less brain activation to complete inhibitory control tasks. Any of the features of complex exercises such as open skills, various motions, and social interaction or those interactions could facilitate neural efficiency. Future studies should investigate what cognitive features of complex exercises induce neural efficiency.

There are several limitations to this study. First, fNIRS measures are easily affected by different experimental protocols. Previous results [28–32] found a significant association between enhanced inhibitory control and increased fNIRS measures by an event-related design for cognitive tasks, not block design as we employed in this study. Compared with event-related design, while block design has the advantage of easily detecting the changes in hemodynamics to cognitive tasks, the disadvantage of block design is easy to contaminate the learning or fatigue effect. Furthermore, types of experimental design might affect the occurrence of neural efficiency. When the difficulty of a cognitive task is low to moderate, neural efficiency in which behavioral performance is improved without increased brain activation trends occurs [43]. However, a challenging cognitive task increases brain activation and results in a correlation between brain activation and cognitive performance [43]. Thus, if the difficulty of cognitive tasks is also affected depending on the types of experimental design, the neural efficiency might not occur when employing an event-related design that makes cognitive tasks challenging. Previous fNIRS studies [49, 50] that employed block design have also reported neural efficiency. The experimental design differences may affect the association between cognitive performance and brain activation. Second, the badminton intervention was not a real match. There was no psychological pressure and stress that badminton players must confront when an actual match. In a real badminton match, psychological pressure and stress may influence inhibitory control and brain activation.

## Conclusions

In conclusion, a single bout of complex exercise selectively enhances inhibitory control relative to a simple exercise. On the other hand, hemodynamics in the prefrontal cortex does not differ in complex and simple exercises. An acute complex exercise could induce neural efficiency after the cession of exercise, enhancing inhibitory control.

## Acknowledgments

The author is grateful to all participants and a badminton instructor.

## Supporting information

S1 File. This is the supporting file data.xlsx.

## Notes

### Competing Interest Statement

The authors have declared no competing interest.

